# Nodulin 16 of *Lotus japonicus* (Nlj16) regulates the recruitment of Leghemoglobin (LegH) to the infected nodule cell membrane during symbiotic nitrogen fixation

**DOI:** 10.1101/2022.07.21.500945

**Authors:** Amit Ghosh, Aniruddho Das, Md Azimuddin Ashrafi, Sudeshna Saha, Firoz Molla, Maitrayee DasGupta, Anirban Siddhanta

## Abstract

Nitrogenase that catalyses anaerobic symbiotic nitrogen fixation (SNF) in legumes is synthesized by rhizobium. Legume root cells express nodulin proteins after infection with rhizobia. Nodulins have been classified as early and late, reflecting the time points of their expression. Leghemoglobin (LegH), which is a classic example of a late nodulin, sequesters oxygen inside the nodule to protect the nitrogenase from oxygen toxicity to sustain SNF. Previous data from our laboratory demonstrated that phosphorylated LegH at S45 showed compromised oxygen sequestration in vitro due to structural disruption of the porphyrin binding pocket responsible for its oxygen binding. Moreover, we have demonstrated by using co-immunoprecipitation that LegH interacts both *in vitro* with Nodulin 16 of *Lotus japonicus* (Nlj16), another late nodulin. Fluorescence Immunohistochemistry (IHC) data shows that both LegH and Nlj16 are localized in the membrane and cytosol of infected cells. Notably, serine phosphorylation of LegH and interaction of Nlj16 with LegH respectively reduces and increases its *in vitro* oxygen sequestration ability. In this report, to further elucidate the spatio-temporal regulation of this interaction, we generated hairy root transgenic *Lotus japonicus* plants where Nlj16 has been knocked down by using siRNA. Most interestingly, our data shows that the membrane localization of LegH is obliterated in the Nlj16 knocked-down root cells infected with the cognate Rhizobia suggesting a distinct role of Nlj16 in its recruitment to the membrane of nodule cells.

## Introduction

Nitrogen fixation is the process by which inert molecular nitrogen is converted to nitrogenous compound naturally or by anthropogenic processes. Biological nitrogen fixation in legumes takes place in their root nodules that result due to the symbiotic interaction between the host plant and endosymbiotic soil bacteria known as Rhizobia. The bacteria convert atmospheric nitrogen to ammonia, in a high energy reaction catalysed by the metallo-enzyme nitrogenase, while the host provides carbon nutrients.

Upon infection, diffusible signalling molecules are mutually produced by legume root cells and the cognate rhizobia. Plant flavonoids and/or phenolics stimulate secretion of Nod factors by the rhizobia which in turn induce expression of set of nodulin genes in the legume root cells (Stougaard 2000; Van De Sande and Bisseling 1997). Nodulin genes have been traditionally classified as early and late, reflecting the developmental time points of their expression. Early nodulin genes that are induced within a few hours of perception of Nod factors are responsible for some important morphogenetic processes such as pre-infection, infection, and cortical cell division etc. Some comprehensive reviews have extensively discussed about the early nodulins (Mylona et al. 1995; Schultze and Kondorosi 1998). After the successful induction of early nodulin genes another set of genes is expressed which is known as late nodulin genes.

Leghemoglobin (LegH), a late nodulin gene product, scavenges oxygen inside the nodule to protect oxygen labile nitrogenase (Downie 2005). On the other hand, Symbiotic Nitrogen Fixation (SNF) requires large number of ATP which is supplied by oxidative phosphorylation in the bacteriod (Kawashima et al. 2001). Therefore, precise maintenance of oxygen microenvironment inside a nodule is very critical to sustain SNF. It is reported in indeterminant nodule of pea plant that, appropriate partial pressure of O_2_ (pO_2_) inside the nodule is partly regulated by both LegH and bacteroid respiration to sustain SNF (Kawashima et al. 2001). Furthermore, affinity of LegH towards oxygen depends on the site of its expression in indeterminate nodules (Kawashima et al. 2001). The exact nature of such regulation is unclear in *L. japonicus* nodules which are determinate in nature where no zone specificity is present.

To address this issue, we have shown that while phosphorylation at specific serine residues of LegH negatively regulates its oxygen sequestration ability, the same is enhanced by the interaction of LegH with another late nodulin, Nodulin 16 of *Lotus japonicus* (Nlj16) *in vitro* (Bhar et al. 2015 and Ghosh et al. 2019). This protein-protein interaction was also shown to occur in 21 days old mature nodule of *L. japonicus* (Ghosh et al. 2019). Here, in this report we have further characterized this interaction *in vivo*.

## Material and Methods

### Plant material and growth conditions

*L. japonicus* (Gifu B-129) seeds purchased from National Bioresource Project (*Lotus japonicus, Glycine max*) University of Miyazaki, Japan. Seeds were used for RNAi experiments. The plants were gown in a controlled environment with a 16-h-day/8-h-night cycle, a 22ºC-day/18ºC-night temperature, and a relative humidity of 70% (Handberg and Stougaard 1992). Before germination, the seeds were surface sterilized for 10 min in conc. H_2_SO_4_ and 10 min in 25% commercial bleach (1% hypochlorite) and 0.1% Triton X, washed 6 times in sterile water and kept in water at room temperature. Then the soaked submerged seeds are transferred to 4°C for 24h. After 24h the seeds are transferred to petri dishes, containing 1% solidified 1/4^th^ B & D nutrient solution (Broughton and Dilworth 1971). The inoculation is performed with *Mesorhizobium loti* (strain NZP2235), and the plants were grown in B & D nutrient solution (Broughton and Dilworth 1971). For studies of nodulation in pots, a sand: soil rite mixture was used (1:2, v/v). Plants were watered three times a week, twice with water and once with nutritive solution, alternately. For nodulation studies N-free nutritive was used.

### Rhizobial strain, growth conditions and inoculation

The *M. loti* wild-type strain NZP2235 (Jarvis et al. 1982) was used for *L. japonicus* nodulation. Rhizobia were grown in YMB for 2 days in the dark at 28°C. *L. japonicus* hairy-root transformed plants were inoculated with *M. loti* suspension at OD 0.01-0.02.

### RNA interference

The 114-bp amplicons were designed against the Nlj16 mRNA sequence from nucleotides 42 to 153. Amplicon specificity was determined through BLAST analysis (https://blast.ncbi.nlm.nih.gov/Blast.cgi?PROGRAM=blastn) against the *L. japonicus* genome. The structure of the RNAi transcripts was predicted using the online software RNAfold and the ‘minimum free energy’ pairing algorithm (rna.tbi.univie.ac.at/cgi-bin/RNAfold.cgi). Primers used for the RNAi strategy are described as follows:

### Primers for RNAi Construct

Forward Primer:

5‘ CAC CAT GAA GAT CTT GCA GCT TGT AG 3’

Reverse Primer:

5‘ TTG TCA GCC GCT GGA ACA 3’

Amplicons were generated from phage display cDNA library of *L. japonicus* (generous gift from Dr. Jens Stougaard) using Phusion_High-Fidelity DNA Polymerase (NEB, https://www.neb.com/) following the manufacturer’s recommendation. The amplicon is then cloned into the pENTR/D-TOPO vector (Life Technologies, https://www.thermofisher.com/). The amplicon was introduced into between the attR1 and attR2 sites of PUB-GWS-GFP binary vector (derived from pCAMBIA 1300) from pENTR/D-TOPO by Gateway LR recombination (Invitrogen, http://www.invitrogen.com/). The recombinant binary vector was transformed into *Agrobacterium rhizogenes* LBA1334 by standard procedure as described below.

### Transformation of *Agrobacterium rhizogenes* with recombinant PUB-GWS-GFP

Competent *Agrobacterium rhizogenes* LBA1334 strain was made following published procedure (Jyothishwaran et al. 2007). The recombinant binary vector was transformed into the competent *A. rhizogenes* LBA1334 by freeze thaw method as described earlier (Jyothishwaran et al. 2007). The transformed bacteria were selected by spreading over LA plates containing rifampicin (50mg/ml), spectinomycin (100mg/ml) and kanamycin (50mg/ml).

### *Agrobacterium rhizogenes* mediated hairy-root transformation of the recombinant binary vector in *L. japonicus*

*Lotus japonicus* hairy-root transformations were performed as described in Kumagai and Kouchi (2003) and Okamoto et al. (2013) with minor modifications using the *Agrobacterium rhizogenes* LBA1334 strain. Hairy roots were subsequently inoculated with *M. loti* and nodules were observed at 21 dpi.

### Generation of antisera against LegH and Nlj16

Custom-made antisera against Nlj16 were generated in guinea pig (BIONEEDS Preclinical Services, Bangalore, India) using affinity purified His-tagged protein. The anti-leghemoglobin antibody was obtained from Dr. C. P. Vance, University of Minnesota, St. Paul, USA (Cordoba et al. 2003). Additionally, we have also raised antisera against purified His-tagged LegH in rabbit. The generation of antisera against Nlj16 (Guinea pig) and LegH (Rabbit) were carried out following guidelines of respective Institutional Animal Ethics Committee. Both antisera were affinity purified using Protein-A agarose beads.

### Fluorescent immunohistochemistry for root nodule sections

Root and nodule samples were harvested from every alternate day from day 1^st^ till day 21^st^ day post infection (dpi). Microtome sections were done using rotary microtome 615 RM2235 (Leica Microsystems) from *L. japonicus* root and/or nodules. Sections (10 to 12μm) were prepared from nodule tissue embedded in Technovit 7100 resin (Hydroxyethylmethacrylate) (HeraeusKulzer) (Grønlund et al. 2005). Multiple nodules from each time point were taken and at least 20 sections were generated for each nodule. Sections were blocked with immuno-buffer (20 mM Tris HCl pH 8.2, 0.9% NaCl, 0.01% BSA and 0.02% Gelatin) and incubated with the IgG fraction of the respective anti-sera (dilution 1:100 in immune-buffer). Alexa fluor (488 and 568)-conjugated anti-rabbit and anti-Guinea pig secondary antibodies (Invitrogen, Thermo Fisher Scientific) (dilution 1:250 in immuno-buffer) were used. Images from multiple fields of view for each section were taken in confocal laser scanning microscope (Olympus, Model IX81) and Leica TCS SP5 II AOBS confocal 619 laser scanning microscope (Leica Microsystems). Z-series sections were acquired at a step size of 0.34μm comprising of about 40 optical sections. 2-D projections were done for Z-series images using FLUOVIEW software (Olympus). Images were processed with FV-10ASW version 4.1 software and prepared for presentation with using 620 Adobe Photoshop CS6. Representative images for each set wereused to make the figure presented here.

### Total RNA Isolation and cDNA preparation

Total RNA was isolated from 21 dpi root nodule of *L. japonicus* by using Macherey-Nagel™ NucleoSpin™ RNA Plant Kit (Cat.No. 38220090). Following total RNA isolation, Cdna was prepared using Invitrogen™ SuperScript™ III First-Strand Synthesis System (Cat.No. 18080051).

### Real Time PCR

Real Time-PCR analysis of the cDNAs was done with Thermo Scientific™DyNAmo ColorFlash SYBR Green qPCR Kit (Cat. No. F-416L) in presence of primers specific for Nlj16 and LjPP2A, the latter being the internal control. Experimental setup and execution were conducted using an Applied Biosystems ABI 7500 Fast Real-Time PCR system. PCR program: 1 cycle at 95°C for 7 mins, 40 cycle at 95°C for 10 sec and 62°C for 30 sec. Expression data were obtained from three independent biological repetition.

### Primers for Real Time-PCR

Nlj16_RT_Forward Primer:

CTGAGCATATAGAGTTTGTCCC

Nlj16_RT_Reverse Primer:

GGACGTTGTGTCATTCTGAC

Internal Control (LjPP2A)

LjPP2A_RT_Forward Primer:

TGCTCCCTCTGGTTGTAAATG

LjPP2A_RT_Reverse Primer:

ACAGGGACGGATGGTATTCT

Relative expression of respective mRNAs was determined by computing 2^-ΔΔCt^ from the output of the RT-PCR process. Comparative 2^-ΔΔCt^ was used to create histogram of relative mRNA transcript level of Nlj16 in the nodules harvested at 21 dpi from WT and Nlj16^KD^ variants of *Lotus japonicus* was represented.

## Results

### Time dependent expression and co-localization of LegH and Nlj16 in wild type nodule

Results from our laboratory showed that the expression and co-localization of both LegH and Nlj16 proteins in root nodules of *Lotus japonicus* (Ghosh et al. 2019). Notably, both the proteins co-localize in the cell periphery and in the cytosolic region in the nodule cells.

Moreover, co-immunoprecipitation data indicates that these two proteins interact in the nodule lysate. To further verify the peripheral localization of both the proteins, we exploited Calcofluor-white which specifically stains plant cell wall. The images of the calcofluor-white stained samples suggest that the peripheral distribution of the two co-localizing protein is in the vicinity of the cell wall which remained discrete (Fig. 1; Arrow heads denoting the co-localized LegH and Nlj16 while arrows indicate the plant cell wall). Furthermore, we observed that the peripheral distribution of the two co-localizing proteins which is more prominent in the samples of earlier dpi declines gradually with the progress of nodule development as evident in the samples of later dpi (Fig. 1). Conspicuously, the decrease in the peripheral localization of these two proteins is concomitant with their increased cytosolic presence.

**Fig. 1.**
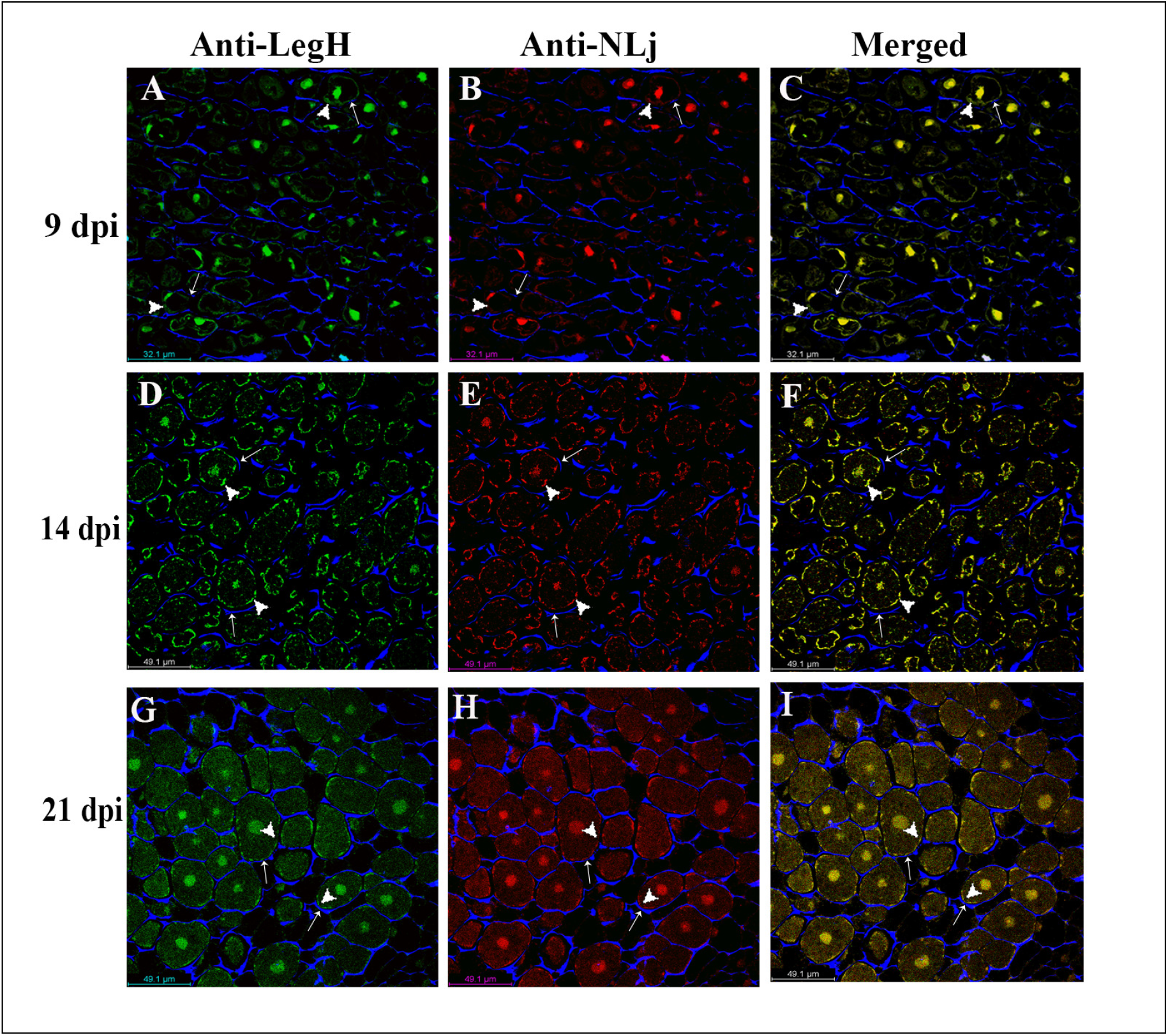
Co-localization of LegH and Nlj16 proteins in the nodules of different dpi in *L. japonicus* by Fluorescent Immunohistochemistry (FIHC). Post-infection nodules were harvested as indicated and processed for IHC. Representative images of the samples from indicated dpi are shown here. Calcofluor-white was used to stain cell walls and the images are shown in blue. The images from red (Nlj16), green (LegH) and blue (Calcofluor) channels were digitally merged and are shown in ‘Merged’ panel. Arrow head shows the complete co-localization of LegH and Nlj16 towards the cellular periphery (most likely cell membrane) while the arrow indicates the position of the plant cell wall in the infected cells. Bars indicate 49.1 μm

### Peripheral localization of LegH is obliterated in Nlj16 Knockdown (KD) nodules

It was demonstrated that Nlj16 and Nlj16 like proteins localize to heterologous plant and yeast cell membranes (Kapranov et al. 2001 and Ghosh et al. 2015). Since, Nlj16 and LegH were shown to interact with each other *in vivo* (Ghosh et al. 2019) therefore we speculated that Nlj16 has a role in the cell surface localization of LegH in the nodule cells. To address this hypothesis, we created transgenic plants of *Lotus japonicus* where Nlj16 was knocked down (KD) using siRNA containing gateway compatible binary vector system. The transcript and protein level of Nlj16 were analysed using RT-PCR and FIHC respectively (Fig 2; A & B respectively). Results for RT-PCR shows ∼80-fold decrease in the mRNA transcript level of Nlj16 in Nlj16^KD^ as compared to that of WT (Fig. 2A). As expected the FIHC result shows that the protein level of Nlj16 was markedly diminished in the nodules form Nlj16^KD^ plants (Fig. 2B). Most interestingly, FIHC performed on 21 dpi nodule of Nlj16^KD^ showed significant disruption in the peripheral localization pattern of LegH in the nodule cells (Fig. 3) as compared to that of vector control (Fig. 3; arrowheads). The cytoplasmic distribution of LegH in these nodule cells remained apparently uninterrupted (Fig 3).

**Fig. 2.**
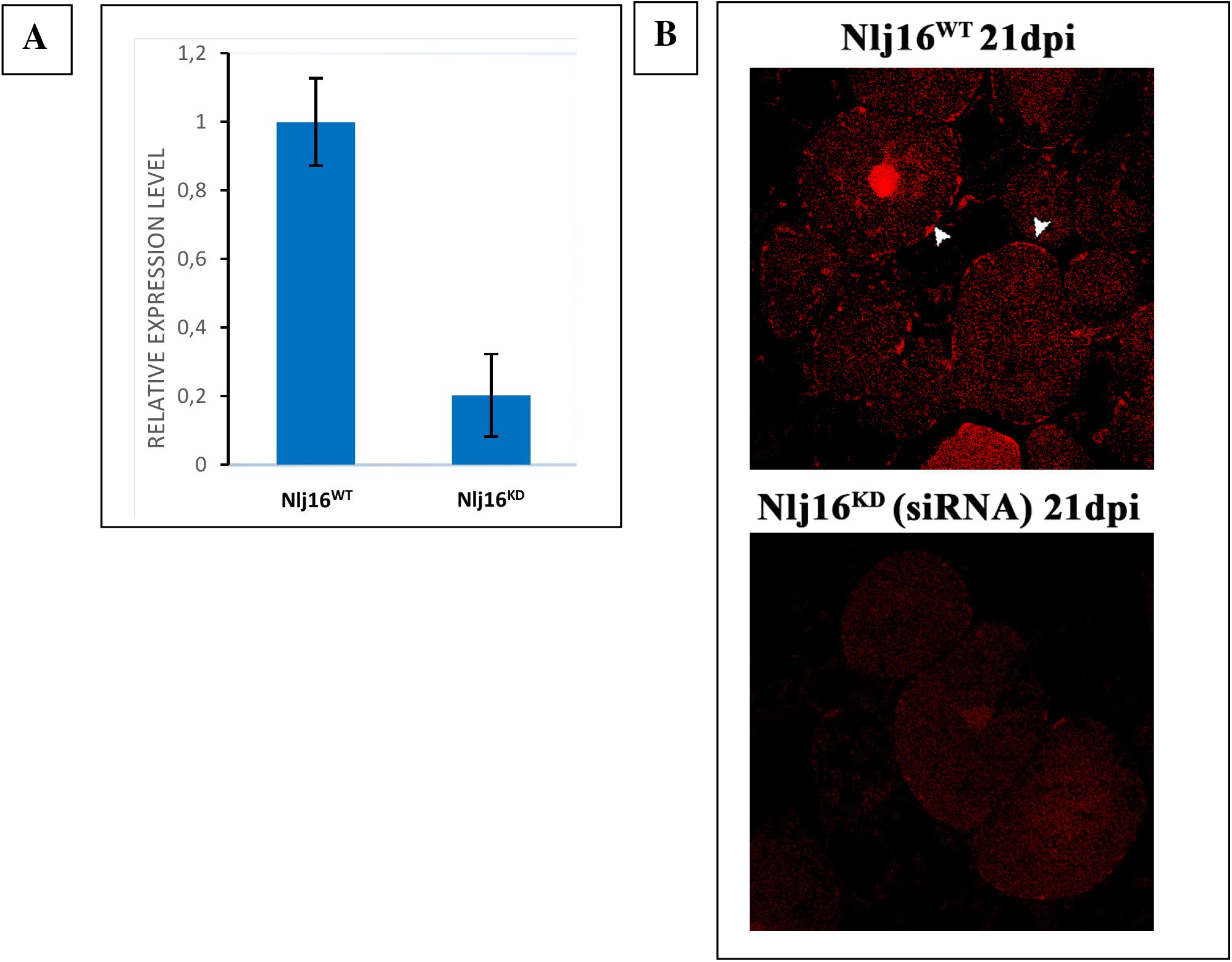
**A. Relative mRNA expression levels of Nlj16 in the nodules harvested at 21 dpi from WT and Nlj16^KD^ variants of *Lotus japonicus*.** Specific primers were used to amplify designated regions of Nlj16 and LjPP2A (as the internal control) from the total cDNA made from respective nodule samples by RT-PCR system. Histogram indicates ∼80-fold decrease in the mRNA transcript level in Nlj16^KD^ as compared to the WT. Error bars represent the standard deviations of the three independent experiments. **B. Reduced Nlj16 protein expression in the nodule sections produced from Nlj16** ^**KD**^. 21-dpi nodules were harvested from WT and Nlj16^KD^ *L. japonicus* plants and processed for IHC as before. Representative images of red (Nlj16) channel of the samples as indicated are shown here.

**Fig 3:**
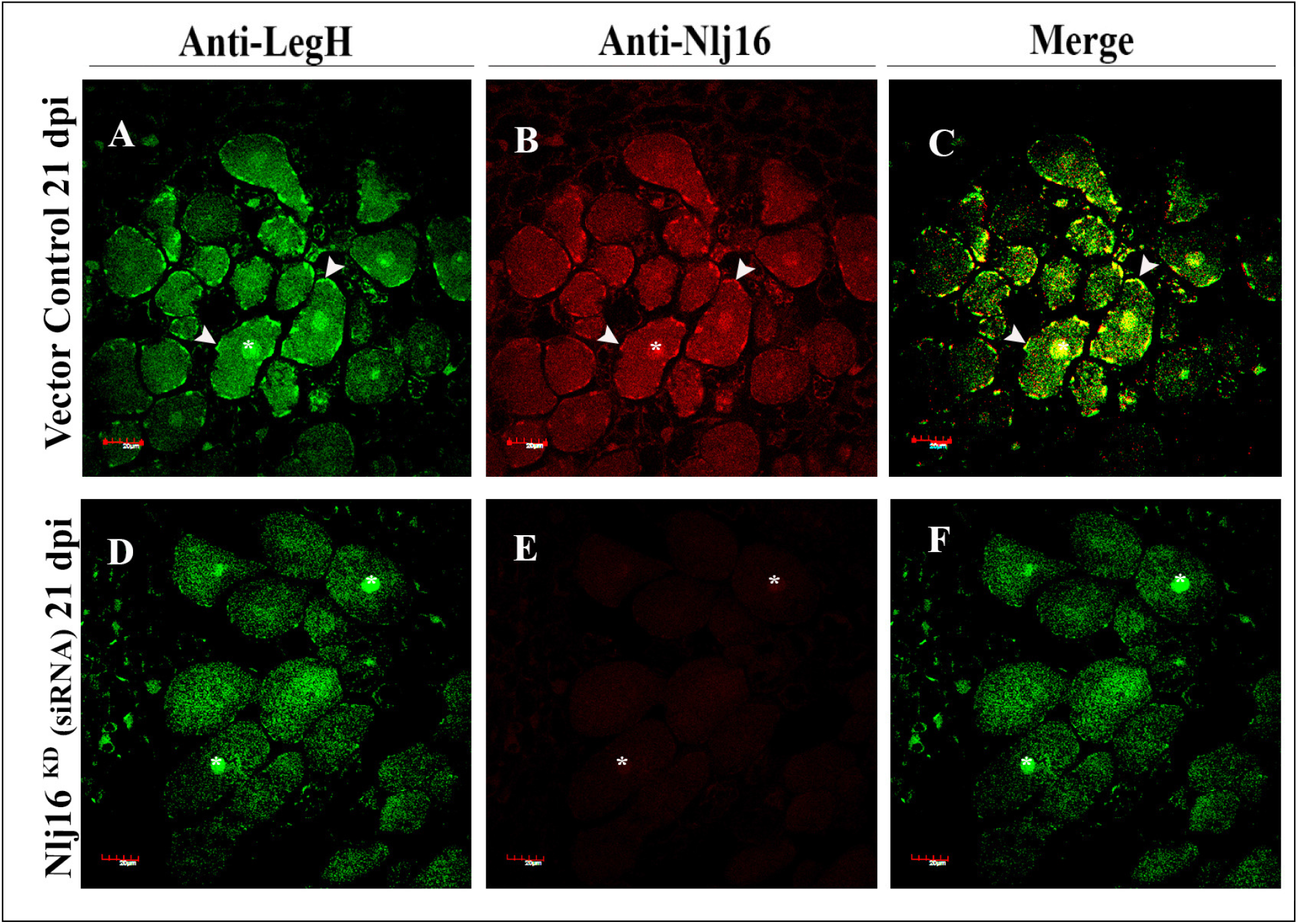
Loss of peripheral distribution pattern of LegH in the nodule cells obtained from the Nlj16^KD^ *L. japonicus* plants. 21-dpi nodules were harvested from WT and Nlj16^KD^ *L. japonicus* plants and processed for IHC as before. The images from red (Nlj16) and green (LegH) channels are shown here. Arrowhead denotes co-localization of LegH with Nlj16 proteins located at the cellular periphery in the panels (A-C). Nuclei of the infected cells are indicated by asterisks (panels A-F).

## Discussion

It is reported that both LegH and bacteroid respiration are responsible to sustain oxidative phosphorylation and SNF (Kawashima et al. 2001). It is known that in indeterminate nodule the LegH is differentially regulated in a zone dependent manner (Kawashima et al. 2001). However, in determinate nodule (such as *L. japonicus*) where no zone specificity is present, the mode of regulation of LegH is not very clear. It is highly likely that LegH might also be temporally regulated in *L. japonicus* nodules as seen in case of *Glycine max* nodules both of which are determinate type (Fuchsman and Appleby 1979).

Results from our laboratory have shown for the first time that *in vitro* serine phosphorylation of LegH negatively regulates its oxygen consumption (Bhar et al. 2015). An existence of conserved serine phosphorylation across the evolutionary border suggests that this phosphorylation of LegH might have a crucial regulatory role during SNF (Lin et al. 2018). Furthermore, we have also shown a novel interaction between LegH and Nlj16 *in vitro* and *in vivo* (Ghosh et al. 2019). This interaction unveils yet another unexplored regulatory aspect of the oxygen sequestration ability of LegH.

We have demonstrated the presence of both LegH and Nlj16 in the cell periphery and cytosol of the infected legume root cells. Notably, the cell surface localization of both the proteins is prominent in samples of earlier dpi with an increased cytosolic distribution in the mature nodule sections where their peripheral presence is decreased (Fig.1). Earlier, it had been shown that the sub-cellular distribution of LegH to be prominent in the cytoplasmic region and within nucleus, while the cell wall, mitochondria, plastid and peribacteriod space shows no specific staining by immunogold staining in pea plant, but the possible reason for such characteristic distribution was unclear (Robertson et al. 1984). Our results also show cytoplasmic and peripheral (adjacent to cell wall) localization of LegH in nodule sections (Fig.1).

Presence of highly expressed late nodulin, Nlj16 in infected nodules of *Lotus japonicus* was detected while screening cDNA library (Kapranov et al. 1997). Primary sequence analyses and other experimental studies revealed that although Nlj16 is a cytosolic protein but also it has the ability to localize in the membrane (Kapranov et al. 2001). Nlj16 like domain was later found to be reported in various proteins like LjPLPs (Kapranov et al. 1997), AtSFH/COW1 (Böhme et al. 2004). It was noted in *Arabidopsis thaliana* that Nlj16 like C-terminal domain present in AtSFH (*<u>A</u>rabidopsis thaliana* <u>S</u>ec <u>F</u>ourteen <u>H</u>omologues) is responsible for cell membrane localization of the protein for its functioning, like root tip growth (Ghosh et al. 2015). Peripheral localization of Nlj16 in our case validates those earlier observations. Here, we hypothesized that the interaction of these two proteins plays a role in their cell surface localization (plausibly cell membrane). Therefore, we thought if that is true then knocking down of one of the proteins would disrupt the localization of the other one. It is obvious that knocking down of LegH would result in lack of SNF in the nodules. Thus, we have knocked down Nlj16 (Nlj16^KD^) using siRNA in a hairy-root transformation system. The nodules sections from Nlj16^KD^ plants clearly showed absence of peripheral localization of LegH leaving behind the cytosolic ones apparently unperturbed in mature nodules (Fig. 3, D-F).

We speculate that the recruitment of LegH by Nlj16 in cell membrane act as a chemical barrier to regulate the entry of oxygen. Our observation partly justifies recruitment of LegH to the cell periphery. Further experiments are in progress to elucidate the regulatory aspects of such localization pattern of these two proteins during SNF.

## Accession number

Nucleotide sequence data for the genes used in this article can be found under the following accession numbers: Nlj16 (LjNOD16), U64964.1 (GenBank); LjPP2A, LotjaGi2g1v0210500.4 (Lotus Base)

## Acknowledgements

This work is funded by Grants from Govt. of India: DST (SERB EMR/2017/004234). Md. Azimuddin Ashrafi, Aniruddho Das and Sudeshna Saha are supported by research fellowships from the University Grants Commission (UGC), India. This research is also supported by DST-FIST (Department of Biochemistry) and DST-PURSE programs of University of Calcutta, India.

## Author Contribution

The idea and planning of the work were undertaken by AG and AS. The experiments were done by AG, AA, AD and SS. The analyses and interpretation of the results were done by AG, AD, AA and AS. The manuscript was written by AD and AS.

